# High precision magnetoencephalography reveals increased right-inferior frontal gyrus beta power during response conflict

**DOI:** 10.1101/2022.04.25.489434

**Authors:** Pria L. Daniel, James J. Bonaiuto, Sven Bestmann, Adam R. Aron, Simon Little

## Abstract

Flexibility of behavior and the ability to rapidly switch actions is critical for adaptive living in humans. It is well established that the right-inferior frontal gyrus (R-IFG) is recruited during outright action-stopping, relating to increased beta (12-30 Hz) power. Additionally, pre-supplementary motor area (pre-SMA) is plausibly recruited during response conflict/switching, relating to increased theta (4-8 Hz) power. It has been posited that inhibiting incorrect response tendencies is central to motor flexibility. However, it is not known if the commonly reported R-IFG beta signature of response inhibition in action-stopping is also recruited during response conflict, which would suggest overlapping networks for stopping and switching. In the current study, we analyzed high precision magnetoencephalography (hpMEG) data recorded with very high trial numbers (total n > 10,000) from 8 subjects during different levels of response conflict. We hypothesized that a R-IFG-triggered network for response inhibition is domain general and also involved in mediating response conflict. We therefore tested whether R-IFG showed increased beta power dependent on the level of response conflict. We also hypothesized that pre-SMA is an important node in response conflict processing, and tested whether pre-SMA theta power increased for response conflict trials. Using event-related spectral perturbations and linear mixed modeling, we found that both R-IFG beta and pre-SMA theta increased for response conflict trials, with the R-IFG beta increase specific to trials with strong response conflict. This result supports a more generalized role for R-IFG beta in response inhibition, beyond simple stopping behavior towards response switching.

**Significance Statement:** Response inhibition is a core component of cognitive control. Neural mechanisms of response inhibition are typically studied using stopping paradigms. However, there is an unresolved debate regarding whether the response inhibition network is specific to stopping or generalizes to switching between tasks and overcoming conflict between competing response tendencies. Increased beta (12-30 Hz) in R-IFG has historically been interpreted as a marker of successful response inhibition in the stop-signal task. Here, we investigated the presence of this electrophysiological marker of response inhibition specifically during response conflict (switching). We found R-IFG beta power increased for trials with strong response conflict, and not for weak or no response conflict, thereby supporting a generalized role for R-IFG beta in response inhibition and switching.

## Introduction

Sometimes we plan or start to execute an action, but then need to suddenly execute a different action instead. In experiments, this has been called switching, response overriding, or overcoming response conflict; here we use the term response conflict. One notable theory suggests that during motoric response conflict, inhibition of the incorrect response tendency is necessary (see Wiecki & Frank, 2013). This inhibitory control mechanism may then be the same as that of outright action-stopping (e.g., Wessel & Aron, 2017; Wessel et al. 2019), although a subsequent response is not required when stopping. However, it remains an open question to what extent response conflict resolution and stopping rely on shared mechanisms of response inhibition.

Various cortical and subcortical regions are thought to facilitate the control of action during response conflict and stopping. Communication from pre-SMA to the STN of the basal ganglia has been implicated in response conflict, and from right-inferior frontal gyrus (R-IFG) to STN in stopping (for review, see Aron et al., 2016; Hannah & Aron, 2021). Medial prefrontal cortex (mPFC), including pre-SMA, has been implicated during response conflict paradigms using neuroimaging (Garavan et al., 2003; Nachev et al., 2005), brain stimulation (Neubert et al., 2010), electrophysiology (Zavala et al., 2018; Wessel et al., 2019), and single-unit recordings (Isoda & Hikosaka, 2007). A common electrophysiological readout during response conflict is increased theta (4-8 Hz) power in mPFC (Zavala et al., 2018; Wessel et al., 2019) – plausibly originating from pre-SMA – and in STN (Zavala et al., 2018). R-IFG has been implicated in stopping, relating to increased beta (12-30 Hz) power (Swann et al., 2009; Wagner et al., 2018; Schaum et al., 2021), which plausibly reflects communication from R-IFG to STN to stop movement.

The notion that mechanisms of response inhibition are important for overcoming such conflict by inhibiting incorrect response tendencies has been supported by computational models (Wiecki & Frank, 2013) and empirical work (Forstmann et al., 2008a; 2008b; Neubert et al., 2010; Brittain et al., 2012; Wessel et al., 2019). However, scant research has implicated R-IFG in response conflict, despite it being a key node in the putative inhibitory control network. Neubert et al. (2010) found using transcranial magnetic stimulation (TMS) that response conflict increased the inhibitory influence of R-IFG over primary motor (M1) cortical representations of incorrect responses. Using fMRI, Forstmann and colleagues (2008a, 2008b) found that R-IFG activation related to behavioral indices of response inhibition during response conflict, although only for some trials. This suggests R-IFG may be recruited to inhibit incorrect responses during response conflict, but the limited evidence is not straightforward, nor has it been supported by high resolution electrophysiology. Brittain et al. (2012) reported increased STN beta during Stroop conflict, which might suggest beta-band communication from R-IFG to STN is involved in response conflict. However, to date there has not been a direct electrophysiological investigation of the role of R-IFG beta during response conflict, which is a focus of the current study.

We re-analyzed a head-cast, high precision MEG (hpMEG) dataset including a very large number of trials (total n > 10,000) from a small cohort of healthy controls (N = 8) during a random dot kinematogram (RDK) response conflict paradigm (Bonaiuto et al., 2018; Little et al., 2019). We reconstructed source activity in pre-SMA, R-IFG, and left-IFG (L-IFG; R-IFG comparison region). We then utilized event-related spectral perturbations and linear mixed modeling to evaluate power changes associated with response conflict at the single-trial level, following the imperative cue which could be congruent or incongruent with the preparatory cue. We hypothesized that R-IFG beta is recruited for response inhibition during response conflict, and tested whether R-IFG beta power increased on response conflict trials, particularly strong response conflict. We also evaluated for classical mPFC theta activity by testing whether pre-SMA theta power increased on response conflict trials. Lastly, we tested whether changes in R-IFG beta and pre-SMA theta during response conflict related to motor behavior.

## Materials and Methods

### Data

The analyses presented in this paper were performed on a pre-existing hpMEG dataset, collected with subject-specific head-casts to maximize signal-to-noise ratio (SNR) (see Little et al., 2018). The analyses were performed with custom scripts in MATLAB R2020a and RStudio Version 1.2.5001. Raw data were obtained from the Open Science Framework (https://osf.io/eu6nx), and directly from the original data collectors, JB and SL. A full description of the original materials and methods can be found in the original description (Bonaiuto et al., 2018). Brief summaries of key features of the initial data collection and processing are included here, along with more detailed information about the methods used that differ from the original techniques.

### Response conflict paradigm

Subjects were presented with RDKs that predicted upcoming movement cues and varied in level of coherence (high, medium, low). The preparatory RDKs were followed by an imperative cue that indicated a left or right button press using the middle or index finger of the right hand, and were either congruent with the preparatory cue (no response conflict) or incongruent (response conflict) (**Figure 1**). We operationally defined low RDK coherence incongruent trials as having a low strength of response conflict, medium RDK coherence incongruent trials as having a medium strength of response conflict, and high RDK coherence incongruent trials as having a high strength of response conflict. On each trial, subjects responded to an imperative cue using either the index or middle finger of their right hand. Between trials, subjects fixated on a central fixation cross (not depicted in **Figure 1**). Eight subjects completed 1-4 sessions each, for a total of 24 sessions in the dataset. We excluded from our analyses any trials with a response time (RT) of less than 100 ms, resulting in a total of 10,496 trials included across all subjects. Neural signals were analyzed after the imperative cue to index response conflict. We did not analyze neural signals during the RDK period, as this has been previously reported (Bonaiuto et al., 2018; Little et al., 2019).

**Figure 1.**
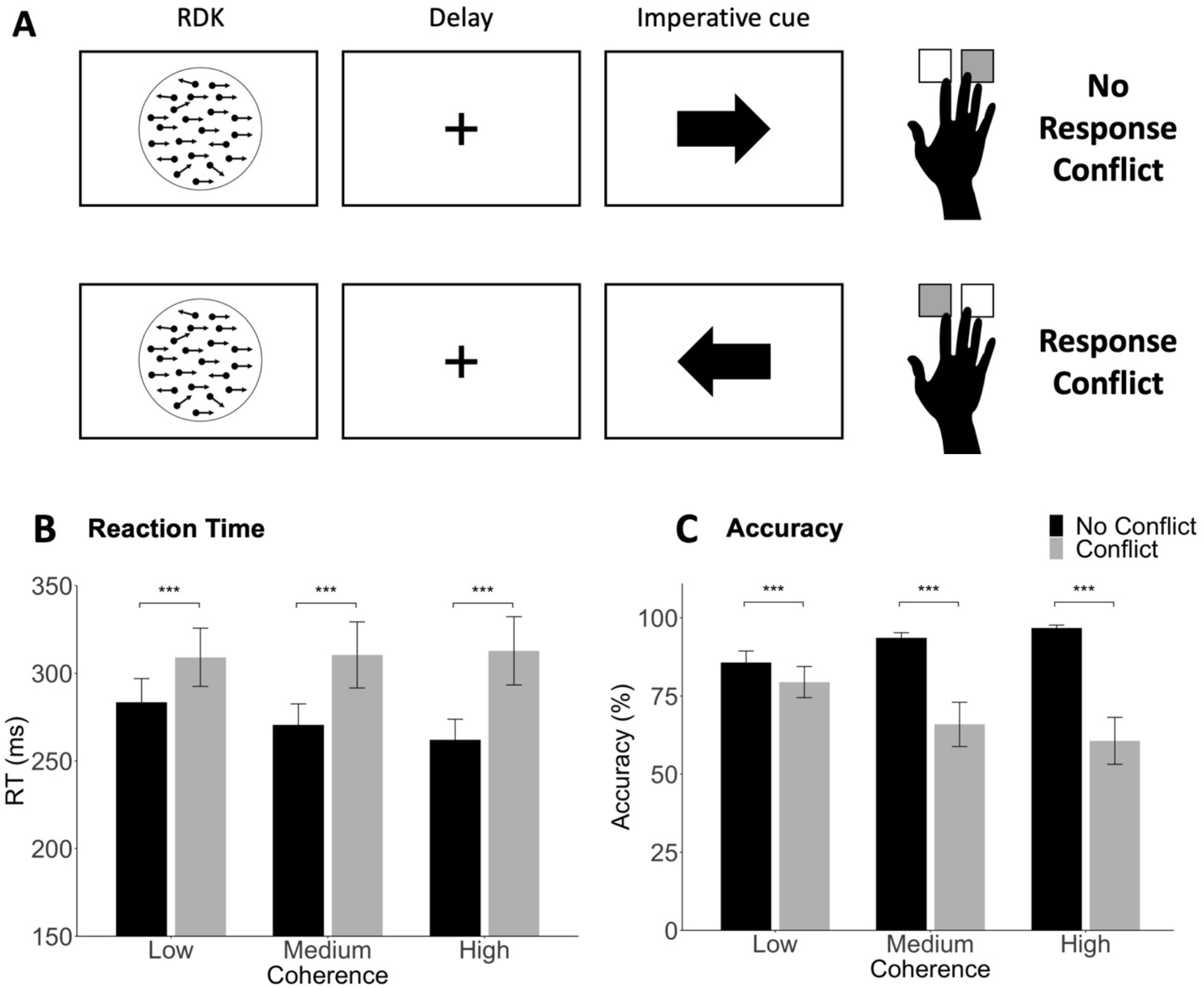
Response conflict paradigm and behavioral performance. **A**. On each trial, subjects were shown a random dot kinematogram (RDK) that varied in level of coherence (high, medium, or low), followed by a fixation cross for a delay period. On a majority of trials (70%), the RDK accurately predicted the direction of the imperative cue, resulting in no response conflict (top panel). On a minority of trials (30%), the RDK inaccurately predicted the direction of the imperative cue, resulting in response conflict (bottom panel). Left and right button press responses were made with the index and middle fingers, respectively, of the right hand. **B-C**. Subjects responded significantly **B)** faster and **C)** more accurately on trials with no response conflict compared to trials with response conflict. Error bars represent standard error of the mean, at the session level (N = 24). *** *p* < .0001.

### Source inversion

Sensor-level hpMEG data (recorded during the response conflict paradigm) was source inverted using individual subjects’ cortical surface meshes to reconstruct activity in pre-SMA, R-IFG, and L-IFG. We analyzed source data from L-IFG for control. For source inversion, cortical surface meshes were extracted using Freesurfer (Fischel, 2012) from multiparameter maps using the proton density (PD) and longitudinal relaxation time (T1) sequences from each subject’s structural MRI, as described by Bonaiuto et al. (2018). 3-dimensional surface plots of each subject’s cortical surface mesh were reviewed and subject-specific regions of interest (ROIs) were established using previously defined anatomical landmarks: right dorsomedial PFC for pre-SMA (see Kim et al. 2010; Neubert et al., 2010; Zhang et al., 2012), and right and left pars opercularis for R-IFG and L-IFG, respectively (see Levy & Wagner, 2011; Breshears et al. 2018). Next, we selected a central vertex within each region, and validated our selection using MNI coordinates from meta-analyses on NeuroSynth (Yarkoni et al., 2011). We created localized clusters by selecting all vertices within a 1 mm radial distance across the cortical surface (see **Figures 3A** and **4A** for a visualization of one subject as an example). We then performed source inversion using SPM 12 (http://www.fil.ion.ucl.ac.uk/spm/) and an Empirical Bayesian beamformer, as described by Little et al. (2019), and then selected source activity time series for every vertex in each ROI cluster for further analysis.

### Time-frequency decomposition

We computed time-frequency transforms of the source-level time series using Morlet wavelets (3 cycles at low frequencies, linearly increasing by 0.5 at higher frequencies), with a range from 4-30 Hz (see Jana et al., 2020). We performed time-frequency transforms for the time series for every vertex in each ROI cluster, to obtain a time x frequency x trial x vertex, 4-dimensional matrix of power for each ROI in each session. Then, we averaged across the cluster vertex dimension to create a time x frequency x trial matrix.

### Event-related spectral perturbation (ERSP)

For group level visualizations, we computed ERSPs for trials with and without response conflict for each ROI (i.e., pre-SMA, R-IFG, L-IFG). We converted spectral power to decibels (dB) using a 500 ms baseline prior to the RDK presentation (i.e., during fixation) (see Cohen, 2014). We averaged across all sessions that each subject completed (total N = 24), and then computed a grand average across subjects (N = 8). We subtracted the group average ERSP for trials with no response conflict from the group average ERSP for trials with response conflict to specifically visualize the difference. We plotted these ERSPs with a range from 4-30 Hz on the y-axis and 0-500 ms relative to the imperative cue on the x-axis, with a vertical line at 300 ms denoting average RT (**Figure 2**). We followed the same procedure for R-IFG ERSPs split by levels of coherence (**Figure 3**).

**Figure 2.**
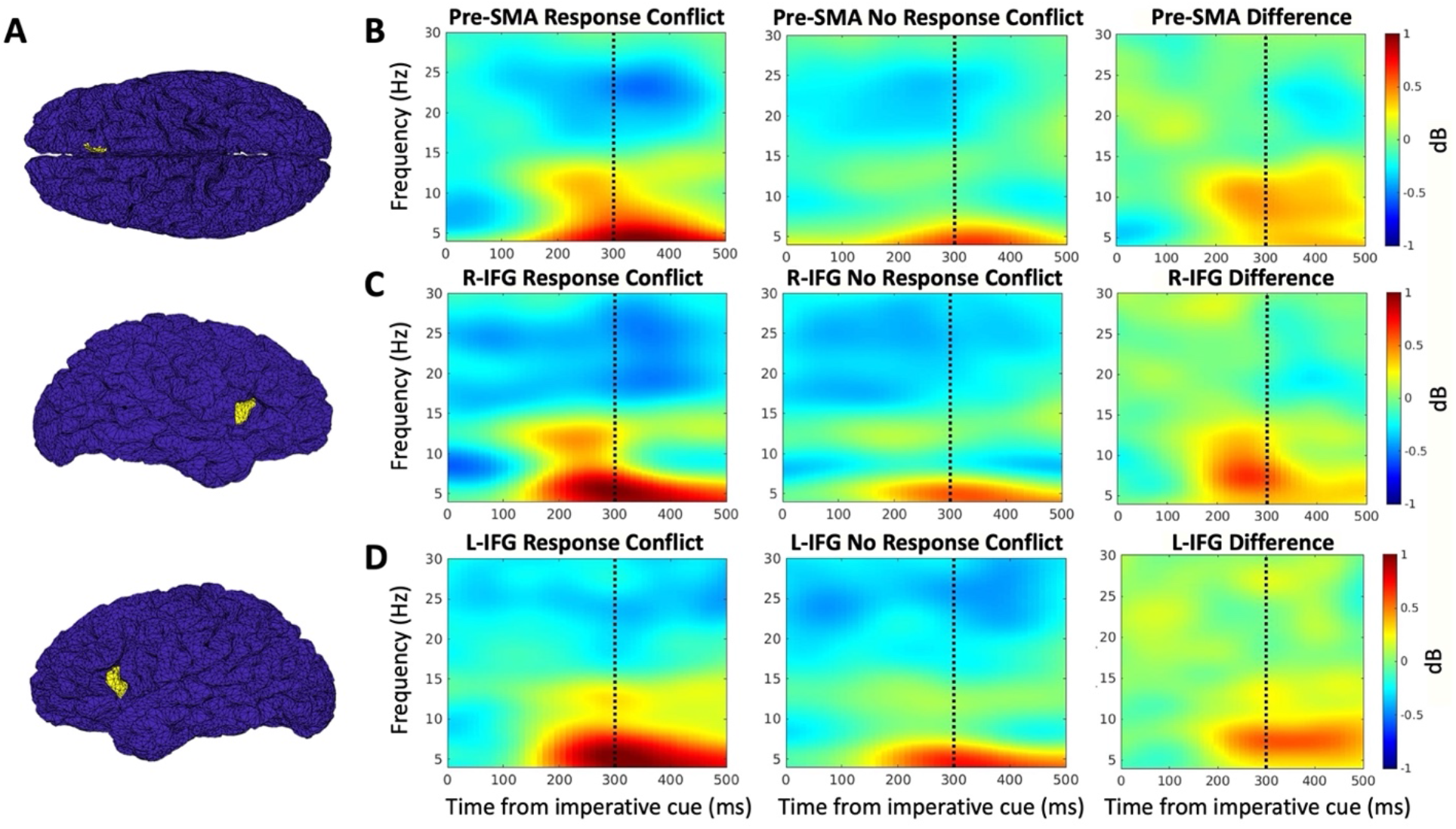
ERSPs for pre-SMA, R-IFG, and L-IFG. Group-average event-related spectral perturbations (ERSPs) for response conflict trials, no response conflict trials, and their difference. Dashed vertical line at 300 ms denotes average RT. Statistics were performed at the single-trial level; these group-level plots are solely for visualization. **A)** Example source localization for pre-SMA (top), R-IFG (middle), and L-IFG (bottom) from one subject. **B)** Pre-SMA shows increased theta and low beta power prior to the average RT for response conflict trials compared to no response conflict trials. Single-trial analyses revealed a marginally significant main effect of response conflict on theta power, and no significant main effects of response conflict on conventional nor subject-specific low beta power. **C)** R-IFG shows increased theta and low beta power prior to the average RT for response conflict trials compared to no response conflict trials. Single-trial analyses revealed significant main effects of response conflict on theta, conventional low beta, and subject-specific low beta power. **D)** L-IFG shows increased theta power (not beta power) prior to the average RT for response conflict trials compared to no response conflict trials. Single-trial analyses revealed a significant main effect of response conflict on theta power, and no significant main effects of response conflict on conventional nor subject-specific low beta power.

**Figure 3.**
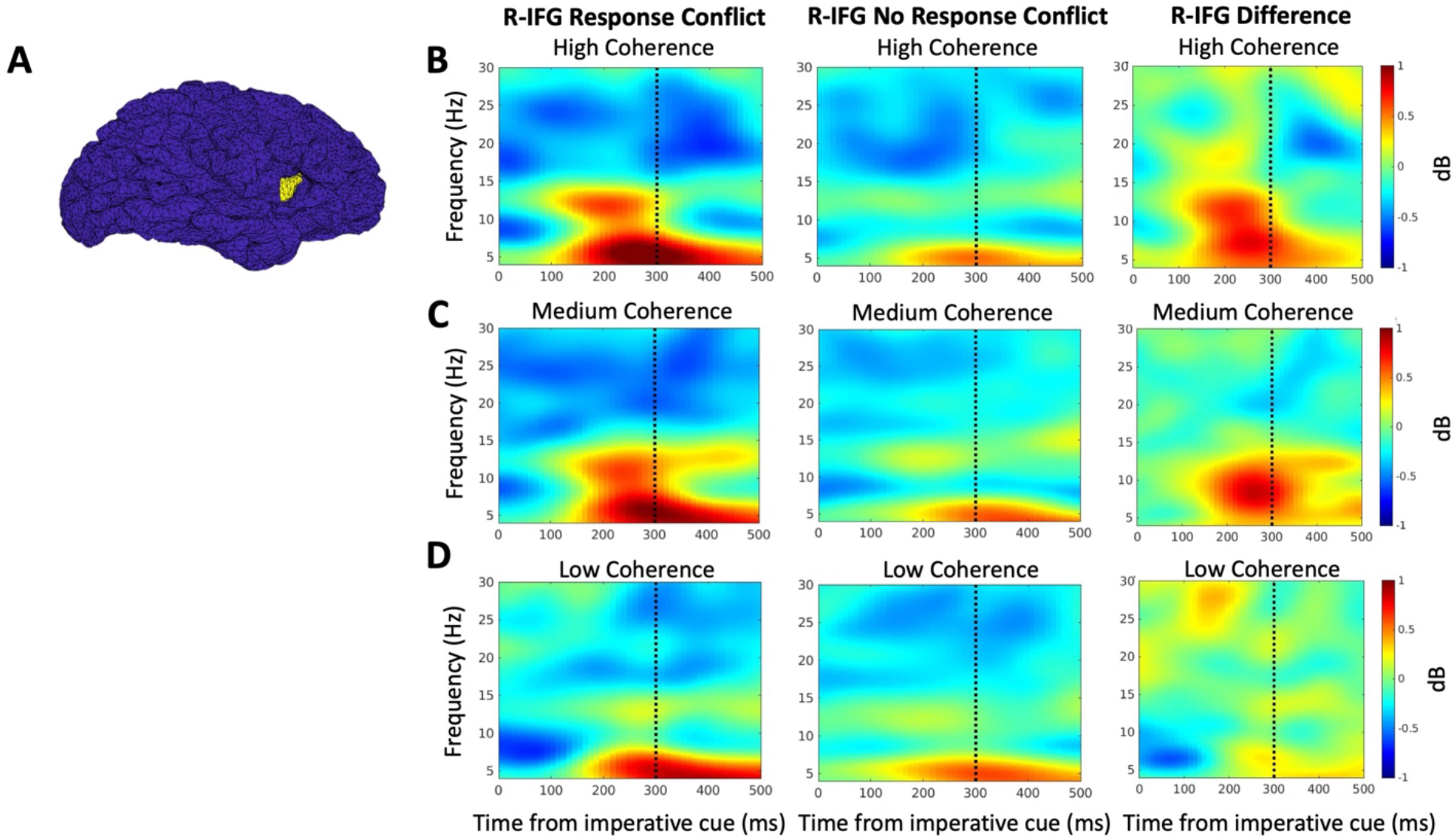
ERSPs for R-IFG, split by coherence level. Group-average event-related spectral perturbations (ERSPs) for response conflict trials, no response conflict trials, and their difference, by coherence level. Dashed vertical line at 300 ms denotes average RT. Statistics were performed at the single-trial level; these group-level plots are solely for visualization. **A)** Example source localization for R-IFG from one subject. **B-D)** R-IFG shows increased low beta power prior to the average RT for response conflict trials compared to no response conflict trials for high coherence (B) and medium coherence (C) trials, and not for low coherence (D) trials. Single-trial analyses revealed a significant interaction between response conflict and coherence on subject-specific low beta power. Pairwise comparisons revealed a significant difference in subject-specific low beta power for response conflict trials compared to no response conflict trials for high coherence trials only, not for medium nor low coherence trials.

### Data preparation for linear mixed modeling

To prepare the spectral power data for single-trial level statistical analyses, we computed a simple linear baseline subtraction using a 500 ms window prior to the RDK presentation on that trial (see Grandchamp & Delorme, 2011; Cohen, 2014). We a priori (prior to visualization of the contrast spectrograms) defined time and frequency ranges to average across to obtain single-trial power estimates. We used 0-300 ms relative to the imperative cue (i.e., time between imperative cue presentation and average RT). We used 4-8 Hz for theta, and subject-specific and conventional partitions of 12-30 Hz for beta (explained in detail below). We z-scored each of these single-trial power estimates within-subject and -ROI.

#### Partitioning the beta band

Various work has defined beta differently (e.g., Engel & Fries, 2010; Newson & Thiagarajan, 2019; Schmidt et al., 2019). Here we defined 12-20 Hz as ‘low beta’ and 21-30 Hz as ‘high beta’. Previous work has specifically implicated the lower beta band in stopping-related activity (Engel & Fries, 2010; Wagner et al., 2018; Schmidt et al., 2019), so we sought as our primary test an evaluation of whether R-IFG low beta specifically was recruited during response conflict. We used an individualized (peak-centered) frequency band of task-relevant low beta to test our primary hypothesis about R-IFG low beta in response conflict. We computed average baseline-corrected spectral power (dB) for all trials that the subject completed, and then averaged across our 0-300 ms time window of interest to obtain a single spectral power estimate for each frequency value in the 12-20 Hz range. Then we extracted the value for which low beta power was greatest, and used that as the center of a narrow subject-specific range (peak low beta +/- 1 Hz). In addition to defining subject-specific low beta, we also used a broader band conventional range of low beta (12-20 Hz) and used the same procedure described here to define subject-specific (peak high beta +/- 1 Hz) and conventional (21-30 Hz) high beta for secondary comparison.

### Experimental design and statistical analyses

#### Linear mixed modeling

To take advantage of the high trial count (n = 10,496) in this dataset, and model within-subject data as well as small cohort between-subject data (N = 8), we used a linear mixed modeling framework using R (v3.6.1, R Core Team, 2019) and the lme4 package (v1.1-21, Bates et al., 2015). All of our models had session nested within subject-specific intercepts as random effects, and the models exploring relationships between neural activity and behavioral performance also had session/subject-specific slopes by response conflict. We used Type III Wald Chi Square tests for our models. We used z-tests, Tukey corrected for multiple comparisons, for pairwise comparisons.

To assess the impact of response conflict and RDK coherence on RT, we used a linear mixed model with RT (log10 transformed) as the dependent variable, and response conflict, RDK coherence, and their interaction as fixed effects. To assess the impact of response conflict and RDK coherence on error, we used a generalized linear mixed model with a binomial distribution, response (0 = incorrect, 1 = correct) as the dependent variable, and response conflict, RDK coherence, and their interaction as fixed effects. To assess the impact of response conflict and RDK coherence on neural activity, we used linear mixed models with z-scored power as the dependent variable, and response conflict, RDK coherence, and their interaction as fixed effects. Lastly, to explore relationships between neural activity and behavioral performance, we used linear mixed models with RT (log10 transformed) as the dependent variable and z-scored power as a fixed effect, and generalized linear mixed models with binomial distributions, response (0 = incorrect, 1 = correct) as the dependent variable, and z-scored power as fixed effects.

## Results

### Behavioral results

On average, subjects responded faster and more accurately on trials with no response conflict compared to trials with response conflict (**Figure 1**). Linear mixed models revealed a significant interaction between response conflict and coherence for log-transformed RT (*X*^*2*^(2) = 54.71, *p* < .0001) and for error (*X*^*2*^(2) = 264.19, *p* < .0001). Pairwise comparisons revealed significant differences in behavioral performance for trials with no response conflict compared to trials with response conflict. Subjects had significantly longer RTs on trials with response conflict for high (*Z* = -19.89, *p* < .0001), medium (*Z* = -16.07, *p* < .0001) and low (*Z* = -9.71, *p* < .0001) coherence trials, and significantly more errors on trials with response conflict for high (*Z* = 26.94, *p* < .0001), medium (*Z* = 20.64, *p* < .0001), and low (*Z* = 4.81, *p* < .0001) coherence trials.

### Neural results

#### Increased theta power during response conflict in pre-SMA, R-IFG, and L-IFG

To test for previously described theta power increases in pre-SMA during response conflict, we analyzed pre-SMA theta power during response conflict trials compared to no response conflict trials. We also analyzed theta power in R-IFG and L-IFG to test for regional specificity of any theta increases. In keeping with the previous literature, theta power in pre-SMA after the presentation of the imperative cue was higher during response conflict trials compared to no response conflict trials, though the effect size of this increase was modest (**Figure 2B**). Our linear mixed model revealed a trend of a main effect of response conflict on pre-SMA theta power (*X*^*2*^(1) = 2.82, *p* = .093). Additionally, theta power in R-IFG and L-IFG after the imperative cue was higher on average during response conflict trials compared to no response conflict trials (**Figures 2C** and **2D**, respectively). Our linear mixed models revealed a significant main effect of response conflict on R-IFG theta power (*X*^*2*^(1) = 12.07, *p* = <.0001) and a trend of a main effect of response conflict on L-IFG theta power (*X*^*2*^(1) = 3.16, *p* = .075).

#### Increased beta power during response conflict in R-IFG, not pre-SMA or L-IFG

To test whether there was increased low beta power in R-IFG during response conflict, we analyzed subject-specific low beta power during response conflict trials compared to no response conflict trials in R-IFG. We also analyzed conventional low beta, and both subject-specific and conventional high beta for comparison. We then analyzed these definitions of beta power in pre-SMA and L-IFG to test for regional specificity of any beta increases. On average, low beta (not high beta) power in R-IFG was higher during response conflict trials compared to no response conflict trials (**Figure 2C**). Our linear mixed models revealed a significant main effect of response conflict on subject-specific low beta power in R-IFG (*X*^*2*^(1) = 4.25, *p* = .039). Our linear mixed models also revealed a significant main effect of response conflict on conventional low beta power in R-IFG (*X*^*2*^(1) = 16.17, *p* < .0001), and no significant main effects of response conflict on conventional nor subject-specific definitions of high beta power in R-IFG.

On average, low beta power in pre-SMA also appeared to be slightly higher during response conflict trials compared to no response conflict trials (**Figure 2B**). However, our linear mixed models revealed no significant main effects of response conflict on subject-specific nor conventional definitions of low nor high beta power in pre-SMA. On average, there appeared to be no differences in L-IFG beta power on response conflict trials compared to no response conflict trials (**Figure 2D**), and our linear mixed models revealed no significant main effects of response conflict on subject-specific nor conventional definitions of low nor high beta power in L-IFG.

#### R-IFG beta power increased for stronger response conflict trials

To test whether the recruitment of R-IFG low beta during response conflict depended on the strength of the response conflict, we looked at the interaction between response conflict and RDK coherence (operationalized as modulating the strength of response conflict on incongruent trials) on subject-specific low beta power, and on conventional low beta power as well. On average, low beta power in R-IFG was higher during response conflict trials compared to no response conflict trials for high coherence trials (**Figure 3B**) and for medium coherence trials (**Figure 3C**), and not for low coherence trials (**Figure 3D**). Our linear mixed models revealed a significant interaction between response conflict and coherence on subject-specific low beta power in R-IFG (*X*^*2*^(2) = 6.76, *p* = .034), and not on conventional low beta power. Pairwise comparisons revealed a significant difference in R-IFG subject-specific low beta power for high coherence (i.e., strong) response conflict trials compared to high coherence no response conflict trials (*Z* = -4.02, *p* = .0001), and not for within medium nor low coherence trials.

### Neural and behavioral results

#### Impact of pre-SMA theta power on RT and error

To test whether the increase in pre-SMA theta during response conflict trials was correlated with behavior, we looked at the main effect of pre-SMA theta power on log-transformed RT and response error. We also looked at whether the increases in R-IFG and L-IFG theta during response conflict related to behavioral performance. Our linear mixed models revealed a significant main effect of pre-SMA theta power on RT (*X*^*2*^(1) = 5.64, *p* = .018), and no significant main effects of R-IFG nor L-IFG theta power on RT. Additionally, our generalized linear mixed models revealed a significant main effect of pre-SMA theta power on error (*X*^*2*^(1) = 8.12, *p* = .0044), as well as significant main effects of R-IFG (*X*^*2*^(1) = 12.39, *p* = .00043) and L-IFG (*X*^*2*^(1) = 11.89, *p* = .00057) theta power on error.

#### Impact of R-IFG low beta power on error for response conflict trials only

To test whether the increase in R-IFG low beta during response conflict trials was correlated with behavior, we first looked at the main effect of R-IFG low beta power on log-transformed RT and on response error, and did the same for L-IFG for comparison. Our linear mixed models revealed no significant main effects of R-IFG nor L-IFG subject-specific nor conventional definitions of low beta power on RT nor error. It is plausible that the relationship between R-IFG low beta power and behavior may only be meaningful on trials with response conflict, if mechanisms of response inhibition are only supporting behavioral performance when there is a need to inhibit an incorrect response tendency. Therefore, we conducted an exploratory analysis restricted to trials with response conflict. Our linear mixed models revealed a trend of a main effect of R-IFG subject-specific low beta power on error (*X*^*2*^(1) = 3.72, *p* = .054), and no significant main effects of L-IFG subject-specific nor conventional beta power on error.

## Discussion

We found support for two theoretically driven, a priori hypotheses about the recruitment of pre-SMA theta and R-IFG low beta during response conflict. We hypothesized that pre-SMA is a critical region for overcoming response conflict, supported by findings in humans and non-human primates (Garavan et al., 2003; Nachev et al., 2005; Isoda & Hikosaka, 2007; Neubert et al., 2010). Specifically, we predicted that pre-SMA theta power increases during response conflict. Electrophysiology studies have shown increased mPFC theta in response conflict paradigms (Zavala et al. 2018; Wessel et al., 2019), which plausibly originates from pre-SMA. This aligns well with a framework implicating theta communication from pre-SMA to STN to M1 to pause motor output until sufficient evidence accumulates to a decision threshold during conflict (see Aron et al., 2016). In the current study we found that, after the imperative cue and prior to the average RT, pre-SMA theta power was increased for trials with response conflict compared to trials with no response conflict, and this increase related to behavioral performance in terms of RT and accuracy in responding. However, the effect size of the theta power increase was modest, with the linear mixed models revealing a trend toward a significant result. We also found that theta increased in R-IFG (significant) and L-IFG (trend towards significance) during response conflict. These latter findings indicate a lack of regional specificity for theta activity which we did not initially predict. This could reflect the high SNR and very high trial number in our data, and could also suggest a broader recruitment of cortical theta during response conflict than has previously been reported.

We also hypothesized that response switching is a generalized form of stopping and therefore would recruit the previously defined R-IFG beta-triggered inhibitory control network during response conflict. A small number of studies have directly implicated R-IFG activity using response conflict paradigms (Forstmann et al., 2008a; 2008b; Neubert et al., 2010). However, these studies have not used high spatial and temporal resolution neuroimaging as afforded by head-cast hpMEG. Increased R-IFG beta activity is usually simply interpreted as a marker of successful response inhibition in the stop-signal task (Swann et al., 2009; Wagner et al., 2018; Schaum et al., 2021). It has been proposed that beta-band communication from R-IFG to STN to M1 stops motor output (see Aron et al., 2016; Hannah et al., 2021), which is supported by increased STN beta power during successful stopping (Ray et al., 2012; Bastin et al., 2014). One notable computational framework (Wiecki & Frank, 2013) posits that mechanisms of response inhibition are important to overcome response conflict, such that one needs to inhibit an incorrect prepotent response tendency in order to execute the correct response in a conflict scenario. This general idea has been supported by various empirical studies (Forstmann et al., 2008a; 2008b; Neubert et al., 2010; Brittain et al., 2012; Wessel et al., 2019), but until now, there has been no direct empirical evidence to support R-IFG beta (i.e., a marker of response inhibition in the stopping literature) during response conflict and action switching.

Importantly, we found here that low beta power in R-IFG was significantly increased on trials with response conflict compared to trials without response conflict. Notably, we did not see any significant increases in beta power in L-IFG. This is a strong comparison region for R-IFG because it is an anatomically matched prefrontal region, but one that is not hypothesized to be a node in the putative inhibitory control network. Another compelling component of this result is that this R-IFG low beta increase during response conflict was specific to higher coherence trials: R-IFG beta power significantly increased for high coherence response conflict trials, quantitatively increased for medium coherence response conflict trials but was not significant, and did not increase for low coherence response conflict trials. We predicted that punctate response inhibition (plausibly via R-IFG beta) might be necessary when the incorrect response tendency is most prepotent (i.e., during strong conflict trials), and our results support this idea. Additionally, we further explored the potential role of R-IFG beta in overcoming response conflict by conducting an exploratory analysis of the main effect of R-IFG low beta power on error for response conflict trials only, and found a trend towards significance. Taken together, these results support a role for R-IFG beta in overcoming response conflict.

Some limitations are worth addressing. We analyzed hpMEG data collected from a small cohort of 8 subjects. Although this is a relatively low number of human subjects, the hpMEG data had a very high SNR and was recorded across a very large number of trials (n = 10,496), similar to primate electrophysiology. Therefore, we were able to perform statistically powerful within-subject analyses that accounted for across subject factors using mixed modeling. This approach was also supported by strong a priori hypotheses that were directly tested here. We also want to be cautious of reverse inference in our interpretation that the presence of R-IFG beta plausibly reflects response inhibition during response conflict. We conclude here that the recruitment of R-IFG beta suggests that mechanisms of response inhibition are at work during response conflict.

This is also supported by the established role of R-IFG beta during action stopping and computational and empirical work suggesting that response inhibition is important for overcoming response conflict (Forstmann et al., 2008a; 2008b; Neubert et al., 2010; Brittain et al., 2012; Wiecki & Frank, 2013; Wessel et al., 2019). Previous electrophysiological work has shown that the right prefrontal beta marker of successful response inhibition during stopping occurs in the lower part of the beta band (Engel & Fries, 2010; Wagner et al., 2018; Schmidt et al., 2019), and our results regarding the significance of low beta, not high beta, are consistent with this. However, it is possible that increased R-IFG beta power during strong response conflict trials here instead reflects a different process. For example, Sharp et al. (2010) implicated R-IFG activity in attention, and response conflict trials were a minority of trials in this paradigm so that may have resulted in subjects’ attention being grabbed on those trials. Nevertheless, Sharp et al.’s (2010) result was not focused on R-IFG beta power specifically, and other studies have shown R-IFG beta as a marker of successful response inhibition in stopping above and beyond differences in attention by using the stop-signal task with attentional-capture trials (see Schaum et al., 2021).

In conclusion, we analyzed hpMEG data with high spatial and temporal resolution recorded during a response conflict paradigm, and found support for two theoretically driven hypotheses about the recruitment of pre-SMA theta and R-IFG beta during response conflict. In a novel result, we showed that R-IFG low beta power was significantly increased for response conflict trials, and specifically strong response conflict trials which plausibly require mechanisms of punctate response inhibition for correct responding. It is plausible that beta-band communication from R-IFG to STN to M1 is recruited to inhibit incorrect response tendencies during response conflict. Future work using causal methods such as TMS or neurofeedback can further establish the role of R-IFG beta power in response conflict resolution. Additionally, other response conflict paradigms, particularly those with high ecological validity, can further investigate the relationship between R-IFG beta activity and behavior, as our marginally significant exploratory result suggests that increased R-IFG beta during response conflict trials may relate to more accurate responding. Overall, our results support R-IFG beta as a neural mechanism of overcoming response conflict, in addition to action-stopping. This broadens the role for R-IFG beta as a domain general inhibitory control signal, which may have clinical implications for populations with inhibitory control deficits.

## Acknowledgments

We acknowledge funding from the Wellcome Trust (105804/Z/14/Z and 203147/Z/16/Z), the Biotechnology and Biological Sciences Research Council (BB/M00965/1), National Institutes of Health (DA026452, NS106822, K23NS120037), and the European Research Council under the European Union’s Horizon 2020 research and innovation programme (ERC-CoG 864550). We also acknowledge Dr Philip Starr for his anatomical expertise in localizing R- and L-IFG regions, Dr Vignesh Muralidharan for support in writing scripts, and Dr Gareth Barnes for supervising the initial data collection.

